# Dynamics of murine brain protein synthesis *in vivo* identify the hippocampus, cortex and cerebellum as highly active metabolic sites

**DOI:** 10.1101/643783

**Authors:** Ser Sue Ng, Jung Eun Park, Wei Meng, Christopher Li-Hsian Chen, Raj N. Kalaria, Neil E. McCarthy, Siu Kwan Sze

## Abstract

Identification of proteins that are synthesized *de novo* in response to specific microenvironmental cues is critical to understanding the molecular mechanisms that underpin key physiological processes and pathologies. Here we report that a brief period of pulsed SILAC diet (Stable Isotope Labelling by Amino acids in Cell culture) enables determination of biological functions corresponding to actively translating proteins in the mouse brain. Our data demonstrate that the hippocampus, cortex and cerebellum are highly active sites of protein synthesis, rapidly expressing key mediators of nutrient sensing and lipid metabolism, as well as critical regulators of synaptic function, axon guidance, and circadian entrainment. Together, these findings confirm that protein metabolic activity varies significantly between brain regions *in vivo* and indicate that pSILAC-based approaches can identify specific anatomical sites and biological pathways likely to be suitable for drug targeting in neurodegenerative disorders.

**Abbreviations:** ApoA1: Apolipoprotein A1, ApoA4: Apolipoprotein A4, ApoE: Apolipoprotein E, ApoJ/Clu: Apolipoprotein J/Clusterin, App: Amyloid-β precursor/A4 protein: App, HDL: high density lipoprotein, Lrp1: Low density lipoprotein receptor-related protein 1, pSILAC: pulsed SILAC, pSIVOM: pulsed-SILAC in vivo labelling in mouse, SILAC: Stable Isotope Labelling by Amino acids in Cell culture)

## Introduction

Rapid changes in cellular proteome are required to support key biological processes including proliferation, differentiation, and migration. Under normal physiological conditions, rates of protein synthesis, modification and degradation are tightly regulated and vary significantly between the cell type, tissue, and molecule in question. In contrast, disruption of normal proteome dynamics is thought to be a key component of the aging process, as well as being implicated in abnormal tissue development(Medina-Cano *et al.*, 2018), and diseases(Steklov *et al.*, 2018) such as inflammation(Yin *et al.*, 2019), Alzheimer disease(Xia, 2019), skeleton dysplasia, cancer and fibrosis(Stegen *et al.*, 2019). However, our current understanding of how protein expression patterns are modified over time and how this varies between different body compartments is extremely limited.

In the brain, regulation of spatiotemporal change in the proteome is critically required for a wide range of normal biological functions including neuron growth, learning and memory formation (Sutton and Schuman, 2006, McClatchy *et al.*, 2007, Gold, 2008, Hernandez and Abel, 2008, McClatchy *et al.*, 2012, Jarome and Helmstetter, 2014). Previous studies have attempted to assess brain protein turnover *in vitro* using a novel ‘stable isotope labelling by amino acids in cell culture’ (SILAC) approach, conducted in parallel with various *in vivo* experiments (Price *et al.*, 2010, Cohen *et al.*, 2013, Dörrbaum *et al.*, 2018). Using an innovative approach of feeding mice with N^15^-labelled blue-green algae prior to conducting mass spec analysis of brain tissues, Pierce *et al*. were able to determine turnover rates for 1010 proteins at the whole organ level (Price *et al.*, 2010), while Cohen *et al*. discerned the half-life of 2802 different brain proteins via proteomic analysis of cortical cultures (Cohen *et al.*, 2013), and Dörrbaum *et al.* used mass spectrometry to assess the stability of >5100 proteins in cultures of rat hippocampus tissue (Dörrbaum *et al.*, 2018). These studies have provided important new insight into protein stability in the brain, but it remains unclear how specific cellular signals trigger active changes in protein translation within different brain regions. Indeed, brain cells interact with their local environment and neighbouring cells via a complex network of signalling pathways that can modify patterns of protein translation, modification and degradation, leading to significant effects on overall organ function. *De novo* protein synthesis in response to a specific stimulus thereby allows rapid generation and transduction of appropriate signalling events that directly influence downstream physiological processes. Consequently, current understanding of the molecular events that regulate key organ functions is limited by our inability to profile actively translating proteins and determine their modification status at specific anatomical sites *in vivo*.

In recent years, SILAC-based labelling has emerged as a powerful method of protein labelling and quantification *in vivo* (Kruger *et al.*, 2008, McClatchy *et al.*, 2015), but current protocols require that animals are fed an expensive isotype-tagged diet for at least two generations prior to conducting experiments (Zanivan *et al.*, 2012). Subsequent development of neutron-encoded (NeuCode) stable isotope metabolic labelling later reduced the minimum feeding period to just 3-4 weeks before investigators were able to perform multiplex analyses of proteome dynamics. Both lysyl endopeptidase (Lys-C) and high resolution MS1 spectra (≥240,000 resolving power @ m/z 400) are required in NeuCode labelling protocol. NeuCode labelling protocol assuming the equal label incorporated efficiency across all isotopologues and taking advantage of direct comparing labelled-to-labelled samples does not required 100% incorporation thereby shorter the feeding time (Baughman *et al.*, 2016, Overmyer *et al.*, 2018). In the current report, with same assumption as NeuCode, we describe a new protocol termed ‘pulsed-SILAC in vivo labelling in mouse’ (pSIVOM) that facilitates analysis of actively translating proteins after just 2 days provision of SILAC diet containing ^13^C-L-Lysine. Proteomic profiling of tissues from these animals by LC-MS/MS enabled robust and reproducible identification of 945 actively translating protein groups in healthy brain from C57BL6/J mice. These highly synthesized proteins were predominantly expressed within the hippocampus, cortex and cerebellum, indicating the capacity of our method to yield site-specific data from within an individual organ system. Our study therefore demonstrates that pSIVOM is a cost-effective and time saving new protocol that supports efficient analysis of proteome dynamics within specific tissues, and will shed important new light on how protein expression profiles change in response to specific stimuli encountered at different body sites.

## Results

### Efficiently labelled of mouse brain proteins with just 2 days pulsed feeding with SILAC diet

Since changing patterns of protein synthesis provide important insight into biological function, we developed a new protocol termed ‘pulsed-SILAC *in vivo* labelling in mouse’ (pSIVOM) that facilitates analysis of actively translating proteins after brief provision of a ^13^C-L-Lysine-labelled diet. For this protocol, male C57BL6/J mice were fasted for 16hrs then transferred onto a SILAC diet with free access to drinking water for 48h thereafter (Supplementary Fig. S1A). Mouse body weight and food consumption were monitored daily (Supplementary Table S1B), and average SILAC diet intake was determined as 0.15±0.01g per mouse per day (Supplementary Table S1 and Fig. S1B). After SILAC pulsing, the mice were euthanized and brain tissues were excised for total protein extraction and proteomics sample preparation either by in-solution digestion (samples Brain1, Brain2, and Brain3) followed with HPLC tryptic peptide fractionation or protein fractionation using SDS-PAGE and in-gel trypsin digestion (Brain4). The extracted and fractioned peptides obtained were then injected into a Q-Exactive LC-MS/MS system for further analysis.

We identified a total of 7569 protein groups in one or more of the biological samples tested, with 4019 being detected in all 4 mouse brains analysed with false discovery rate (FDR) <1% for both peptides and proteins, minimum unique high confident peptide threshold set to 1). Within these datasets, average ^13^C-L-Lysine incorporation was 4.7% at the peptide group level and 23.5% at the protein group level (Table 1), with 1890 individual protein groups being heavy-labelled across all four data sets, representing 432 quantifiable protein groups (Supplementary Tables S2 and S3). In these groups, heavy/light chain abundance ratio (after normalization with all peptides) ranged from 0.055-100, with correlation values 0.72-0.94 between biological replicates (Supplementary Table S3 and Fig. 2A-F). These data confirmed that pSIVOM supports efficient identification of brain proteins that are being actively translated *in vivo* after only a short duration SILAC diet.

**Table 1.**
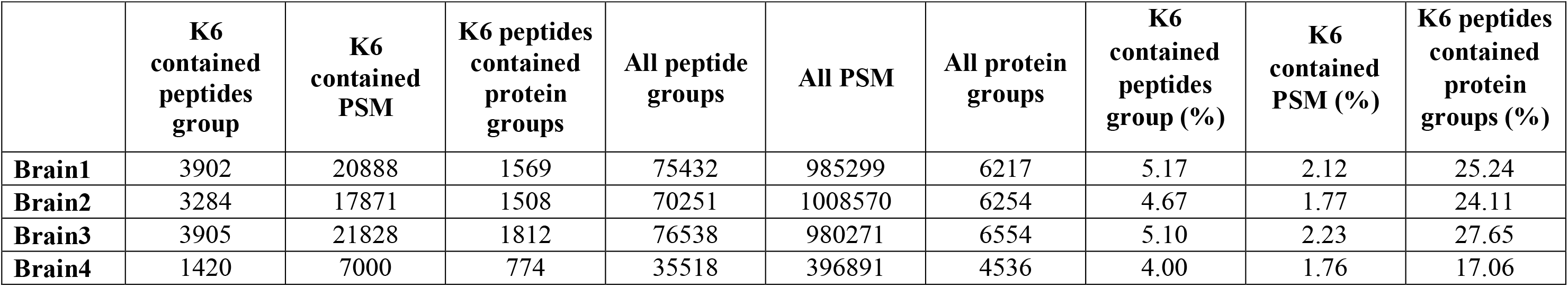
Murine brain-derived peptide spectrum match (PSM), peptide and protein groups as identified within biological samples Brain1, Brain2, Brain3 and Brain4.

### Hippocampus, Cortex and Cerebellum are the most active protein translating region in mouse brain

We next focused on datasets corresponding to brain samples 1-3 which had each been processed using an identical protocol. In this subgroup, we identified a total of 5464 protein groups across all three datasets, including 945 distinct groups found to be both heavy-labelled and quantifiable with biological correlation >0.86 (FDR <1% for both peptides and proteins, minimum unique peptide threshold set to 1). We then further narrow down our candidate list by additional filtering factor, “Abundances grouped standard error in percent”. Total of 823 protein groups were selected with both light and heavy abundance grouped standard error <40%. These candidate protein groups’ biological correlations are >0.93 (Table 2 and Fig. 3A-D) and their technical replicate correlations are all ≥0.90 (Supplementary Table S4, Fig. S2). As shown in Fig. 4, we found that these 823 protein groups’ average heavy/light abundance ratio(log2) were distributed between −0.5 to 1, suggesting moderate *de novo* expression within the tissue of origin. We then performed DAVID gene clustering analysis on these 823 protein groups (or 808 genes) (EASE cut-off value 0.01), which revealed that the newly synthesized proteins were predominantly expressed in the hippocampus (32.7%), cortex (14.9%), and cerebellum (12.9%) (Table 3A). Based on mean values, protein groups clustering in the visual cortex and Corpora quadrigemina were found to be more actively produced than those associated with other regions (Fig. 5, Table 3B). When further assessed according to cellular composition and/or reported function, the actively translating proteins were found to be are mainly located in the cytoplasm (65.8%) followed by the outer membrane (53.0%), and extracellular exosomes (51.4%), with only a minority conferring to nuclear contents (40.5%) (Table 4A). Crucially, several protein subsets were observed to be directly related to key brain cell functions, including myelin sheath biology (13.8%) and neuron projection (11.4%), as well as components of the neuronal cell body (10.9%), axons (9.2%) and dendrites (8.9%). Reflecting the highly dynamic proteomic regulation of these brain cell compartments, the top 5 functions identified among heavy-labelled proteins were biomolecule transport, phosphorylation, oxidation-reduction events, cell-cell adhesion/binding, and protein translation (Table 4B and C). Both the biological processes and molecular functions identified by these clustering analyses suggested that actively synthesized brain proteins are primarily mediators of biomolecule transport and translation.

**Table 2.** ^13^C-L-Lysine labelled protein groups identified with high confidence in brain tissue samples 1-3. The correlation between biological triplicates are indicated.

**Table 3.** Hippocampus, brain cortex and cerebellum are the most active translating regions in brain. (A) DAVID cluster analysis of tissue expression for selected protein groups (EASE cut-off value 0.01) (B). Protein expression and average H/L abundance ratio (Log2) in specific brain regions.(C) Selected candidates and their uniprot accession ID used during DAVID clustering searches and their average H/L abundance ratio [Log2] between the biological triplicates.

**Table 4.** DAVID clustering search results on selected 808 genes. (A) cellular compartment (B)biological process (C) molecular functions and (D) KEGG pathway results (EASE cut-off value 0.01) on selected genes. (E) Gene mapping list for selected genes in the SynSysNet synaptic database. Candidates that found in SynSysNet synaptic database are high-lighted in red.

### Metabolic related protein and Synaptic functional related proteins are the most heavy labelled protein clusters

Consistent with the fact that the brain exhibits the highest energy consumption rates in the body, the majority of newly synthesized proteins detected were found to be involved in various metabolic pathways (Table 4D). We also detected active expression of proteins known to participate in various synaptic processes including signalling via the dopaminergic / cholinergic / glutamatergic / GABAergic pathways, as well as vesicle cycling, long-term potential, short term depression, and the cGMP-PKG (cGMP-dependent protein kinase or Protein Kinase G) cascade. Indeed, when these findings were compared with the synaptic protein database provided on SynSysNet, we were able to confirm that 322 out of these 808 genes (39.8%) were matched with the genes provided (Table 4E). SynSysNet (http://bioinformatics.charite.de/synsys/) is “A European expertise Network on building the synapse” and providing downloadable synaptic protein database. Their database contains 1028 genes that adequate defined pre- and post-synaptic proteins as well as proteins present in sub-domains of the synapse such as synaptic vesicle and associated proteins, lipid rafts and postsynaptic density. This indicating 31.3% synaptic functional related genes from the database are found to be active translating proteins in our study. Less prominent features of the heavy-labelled proteins were circadian entrainment signalling (2.1%), neurotrophin signalling (2.7%), and sphingolipid / axon guidance signalling (2.5%). Intriguingly, we were also able to identify a protein cluster previously implicated in multiple different brain disorders including Alzheimer’s (4.3%), Huntington’s (4.2%), and Parkinson’s disease (3.6%).

### Brain cholesterol homeostasis key enzymes are found to be highly heavy labelled

In addition to high energy use, the brain is also the most lipid-rich organ in the body (Dietschy, 2009), and employs distinct lipid/lipoprotein metabolic pathways in order to cope with segregation from other tissues behind the impermeable blood-brain barrier (Zlokovic, 2008). Brain lipids consist of glycerophospholipids, sphingolipids and cholesterol (Korade and Kenworthy, 2008), with previous studies suggesting that almost all cholesterol in the CNS(central nervous system) is synthesized *de novo* and exhibits a half-life of 0.5-5 years (compared with just a few days for blood plasma cholesterol). Adult brain is also known to contain ~20-25% of total cholesterol in the body (Bjorkhem and Meaney, 2004, Dietschy, 2009), with majority of this being unesterified content within the oligodendrocytes of myelin sheaths, as well as the plasma membranes of astrocytes and neurons(Snipes and Suter, 1997, Dietschy and Turley, 2004). Steady-state maintenance of constant cholesterol levels in the brain is therefore essential for normal morphology and function, hence disruption of cholesterol homeostasis in this organ is linked to several neurodegenerative disorders including Alzheimer’s, Parkinson’s, Smith-Lemli-Opitz syndrome and Nienman-Pick type C disease(Martin *et al.*, 2014). As showed on Table2, it is interesting to found several heavily labelled enzymes are those crucial for brain cholesterol homeostasis, including (1) cholesterol synthesis: Hmgcs1(3-Hydroxy-3-Methylglutaryl-CoA Synthase 1) which display average heavy/light abundance ratio 7.3±1.1 and emPAI value of 4.1±1.8; (2) cholesterol ester : ACAT1(Acyl-CoA cholesterol acyltransferase1) with average heavy/light abundance ratio 0.5±0, emPAI value 13641±418.7; (3) intercellular cholesterol trafficking: Apoa1, Apoa4, Apoe, ApoJ/Clu, Lrp1(Low density lipoprotein receptor-related protein 1), which will be described more detail later; (4) cholesterol excretion: Cyp46a1 (Cholesterol-24-hydroxylase, average heavy/light abundance ratio 3.9±0.3, emPAI value 5.6±2.4) which is a key enzyme for brain-specific cholesterol export form, the 24-HC (24(S)-hydroxycholesterol) production. Few others essential enzymes for cholesterol homeostasis in brain, such as HMG-CoA (3-hydroxy-3-methylglutarly-coenzyme A), HMG-CoA reductase, DHCR24 (lanosterol-converting enzymes-24-dehudrocholesterol reductase), CYP51 (lanosterol 14-alpha demethulase), ATP-binding cassette transporters(ABCA1, ABCG1, ABCG4) etc, indicating these enzymes are either has much lower turnover rate or the expression is relatively too low for quantification.

## Discussion

### ApoA1, ApoA4, ApoE and ApoJ are the most active translating apolipoprotein in mouse brain

Apos (Apolipoproteins) play pivotal roles in the transport and metabolism of lipids within the CNS, where trafficking is mediated specialized ‘high density lipoprotein (HDL)-like particles’ enriched in ApoE/ApoA1(Demeester *et al.*, 2000, Koch *et al.*, 2001, Balazs *et al.*, 2004). Among the apolipoproteins identified to date, nine of 22 have previously been detected at mRNA and/or protein level in the CNS (*ApoC1, ApoC2, ApoD, ApoE, Clu/ApoJ, ApoL2, ApoL3*, and *ApoA4*)(Elliott *et al.*, 2010). In the current study, we observed active synthesis of ApoA4, ApoA1, ApoE and ApoJ/Clusterin in healthy mouse brain, with each of these molecules displaying biological triplicates average Heavy/Light abundance ratio all >10 (log2 value>12) (Table2). Intriguingly, Apoa4 has previously been suggested to be expressed at lower levels than other apolipoproteins in the Sprague-Dawley rat brain (Liu *et al.*, 2001, Shen *et al.*, 2008), but here we detected higher levels of heavy labelling in this protein than were observed for other family members (average Heavy/Light abundance ratio=28.7±1.5, average emPAI=1.4±0.5, 3 biological replicates). ApoA4 synthesis is typically thought to be confined to the intestine, although low-level expression has also been reported in the hypothalamus or prefrontal cortex (Liu *et al.*, 2001, Shen *et al.*, 2008, Elliott *et al.*, 2010). The primary function of these molecules in lipid metabolism remain somewhat unclear, but roles in satiety and appetite regulation as well as anti-oxidant and anti-atherogenic properties have been identified in rodent models (Okumura *et al.*, 1994, Tso *et al.*, 1995, Duverger *et al.*, 1996, Tso *et al.*, 1999, Ostos *et al.*, 2001, Recalde *et al.*, 2004, Wong *et al.*, 2004). Polymorphisms in the ApoA4 gene have also been reported to enhance activation of LCAT (Lecithin:cholesterol acyltransferase) and potentially increase Alzeimers’ disease risk(Csaszar *et al.*, 1997). Our data now suggest that ApoA4 likely plays an important role in the CNS that depends on active synthesis and rapid degradation to maintain low-level expression in the healthy brain.

Another key member of the apolipoprotein family is ApoA1 which is the major protein constituent of plasma HDL. In addition to high expression in several peripheral tissues including the liver and intestine, ApoA1 is also one of the most abundant apolipoproteins in CSF(cerebrospinal fluid)(Roheim *et al.*, 1979, Pitas *et al.*, 1987, Harr *et al.*, 1996) and also serves as an important cofactor on LCAT activation (Sorci-Thomas *et al.*, 2009, Cooke *et al.*, 2018). Indeed, our data indicated that ApoA1 is far more abundant than ApoA4 in the murine brain (emPAI value 33.8±13.1) while still displaying a high level of heavy labelling by pSIVOM analysis (average heavy/light abundance ratio 16.9±1.5). An earlier study has linked early onset Alzeimers’ disease with a polymorphism (−75A/G) in the promoter region of the ApoA1 gene which conferred a modest increase in plasma levels of this protein, however increase of ApoA1 level in CSF of Alzeimers’ disease and dementia patients are still controversy. (Jeenah *et al.*, 1990, Tuteja *et al.*, 1992, Angotti *et al.*, 1994, Harr *et al.*, 1996, Juo *et al.*, 1999, Demeester *et al.*, 2000, Vollbach *et al.*, 2005). These findings suggest that analysis of proteome dynamics in the brain, and apolipoprotein biology in particular, could lead to new insight into the molecular basis of major neurological disorders. Indeed, our study also detected high CNS expression of ApoE which serves as the major transport protein for extracellular cholesterol and other lipids in this compartment. Since there is known to be no exchange of ApoE between the brain and peripheral pools (Linton *et al.*, 1991), we can be confident that the heavy-labelled protein detected here was newly synthesized locally in the brain. In healthy adult brain, nascent ApoE lipoprotein synthesis occurs mainly in astrocytes, cholesterol is then transferred to ApoE to form a mature lipidated ApoE-containing lipoprotein particle that can be uptake by surrounding neurons via binding to lipoprotein receptors (Elshourbagy *et al.*, 1985, Pitas *et al.*, 1987, Wahrle *et al.*, 2004, Kim *et al.*, 2007). Under physiological conditions, ApoE protein levels are relatively stable and mediate dynamic transfer of lipids between brain cells in the CNS, whereas detrimental events such as injury can lead to dramatic increases in glial/neuronal levels of ApoE up to 150 folds (Ignatius *et al.*, 1986, Snipes *et al.*, 1986). In our study, ApoE displayed similar abundance levels and synthesis rates to ApoA1 (average heavy/light abundance ratio 12.6±1.3, emPAI value 31.1±6.7), confirming that multiple apolipoproteins are rapidly expressed in the murine brain. Indeed, while present at markedly lower levels, the alternative family member ApoJ/Clu also displayed a heavy labelling profile consistent with active synthesis (average heavy/light abundance ratio 10.4±1.4, emPAI value 4.2±0.8). ApoJ/Clu is more widely distributed around the body than ApoA1, ApoA4 and ApoE, being expressed in multiple peripheral organs as well as the brain. Within the CNS, ApoJ is primarily produced by astrocytes, but this protein can also be detected in pyramidal neurons of the hippocampus and Purkinje neurons in the cerebellum(Garden *et al.*, 1991, Pasinetti *et al.*, 1994). Previous studies have identified that neuronal expression of ApoJ/Clu can be significantly upregulated by stresses including cytotoxic insult and cellular injury(Dragunow *et al.*, 1995, Klimaschewski *et al.*, 2001, Iwata *et al.*, 2005). Together, these data confirm that the dynamics and distribution of apolipoprotein expression in the brain are key components of healthy brain function, and that dysregulation of this biology is highly likely to confer disease. Indeed, both ApoE and ApoJ have been identified as genetic risk factors for development of late-onset Alzheimer’s disease due to their crucial role in mediating degradation and clearance of App(Amyloid-β precursor/A4 protein) in the brain(DeMattos *et al.*, 2004, Harold *et al.*, 2009, Lambert *et al.*, 2009). Consistent with these data, App also displayed marked heavy labelling/active synthesis in healthy mouse brain tissues subjected to pSIVOM analysis (Heavy/Light ratio: 10.0±0.3, emPAI value: 3.8±1.0).

### Critical enzymes involved in App proteins processing and stability are fast turnover rate proteins in mouse brain

β- and γ-secretases mediate App cleavage at the amino-terminus and carboxyl-terminus respectively, there by generating the amyloid-intracellular domain(AICD), soluble ectodomains sAPPβ and β-amyloid peptides (Aβ) and enabling extracellular deposition of Aβ, which is a key event in the formation of β-amyloid plaques/senile plaques. Multiple studies have now reported that different isoforms of ApoE (E2, E3 and E4) in human not only exhibit differential binding affinity for App(Strittmatter *et al.*, 1993, Holtzman *et al.*, 2000), but their relative expression levels can also confer increase risk of neurodegenerative disorders and stroke(Slooter *et al.*, 1997, Jha *et al.*, 2008). Expression of a mutated form of human App precursor protein in a transgenic mouse model leads to significant App deposition in the brain, but these deposits are substantially reduced by performing the same experiment in mice with an ApoE knockout background (Berul *et al.*, 1999, Fagan *et al.*, 2002). ApoE4 is the single largest genetic risk factor for sporadic AD and promotes disease pathology by seeding Aβ aggregation in the brain, but recent data indicate that AD risk can be reversed by loss of the neuronal receptor Lrp1 (Tachibana *et al.*, 2019) which was identified in our experiments as undergoing active synthesis in murine brain (average heavy/light ratio: 4.9±0.2, average emPAI value: 1.8±0.2). These data indicate that pSIVOM can provide novel insight into the biology of human ApoE isoforms through study of human ApoEs knock in mouse brain proteomics, which is central to AD pathology in human patients.

Other β-amyloid binding proteins were also prevalent in our datasets, including Apbb1, Apba2 and Itm2b which contribute to App processing and protein stability. Apbb1 (Amyloid beta A4 prevursor protein-binding family B member 1) is an adaptor protein that localized in the nucleus and can interacts with App, low-density lipoprotein receptor and transcription factors. It can form a complex with the γ-secretase-derived App intracellular domain and modulated App turnover and processing(Chow *et al.*, 2015), suggesting a potential role for this protein in the pathogenesis of AD. However, our analyses also identified active synthesis of Apba2/Mint2/X11, which instead stabilizes App and inhibits production of proteolytic fragments including the Aβ peptide that is characteristically deposited in brain tissues from AD patients(Yoon *et al.*, 2007, Saito *et al.*, 2008). Similarly, the C-terminal ectodomain of Itm2b/BRI2 can undergo a series of processing events that ultimately generate the soluble peptide BRI2C, which can inhibit App aggregation and fibril deposition(Matsuda *et al.*, 2005, Kim *et al.*, 2008). These data confirmed that our approach can not only be used to elucidate the molecular basis of neurological diseases, but also the mechanisms that protect against brain pathology. Indeed, proteasome-mediated proteolysis is known to be crucial for synaptic plasticity in both mice and humans (reviewed (Hegde, 2010)), and our analyses identified a wide range of different subunits and interacting partners of this complex (Table2). In particular, proteasome 26S subunit ATPase 4 (Psmc4) displayed high abundance (average emPAI value 40.3 ± 5.9) and was the only protein found to exhibit 100% incorporation of heavy lysine after just 2 days SILAC diet and detected consistently across all 4 biological replicates. These data indicate that Psmc4 has an extremely high turnover rate in healthy brain and suggests that proteasome components may themselves be degraded via the ubiquitin proteasome pathway. The results indicated that these actively translating proteins were critically important in maintaining brain proteostasis, and any perturbation and imbalance in these protein translation can potentially lead to proteinopathy and neurodegeneration.

### Ndrg4, Qki, Rasgrf1, Atxn10 and Nedd4 are highly dynamic proteins involved in neuronal development processes

An additional subset of extensively heavy-labelled proteins in our dataset were related to processes of neuronal development (Table 4B), including Ndrg4, Qki, Rasgrf1, Atxn10 and Nedd4. Cytoplasmic protein Ndrg4 (NDRG4/N-myc downstream-regulated gene 4 protein) is known to contribute to steady-state maintenance of intracerebral BDNF(brain-derived neurotropic factor) levels, which is critical for spatial learning and resistance to neuronal cell death during ischemia (Yamamoto *et al.*, 2011). Qki (KH domain containing RNA binding) is a RNA-binding protein that influences glial cell fate and development(Hardy, 1998), but also plays an essential role in myelinisation processes, since spontaneous mutations in this protein result in hypomyelinization of the central and peripheral nervous systems (Hardy *et al.*, 1996, Lu *et al.*, 2003, Larocque and Richard, 2005). Rasgrf1 (Ras protein-specific gunine nucleotide-releasing factor1) is a guanine nucleotide exchange factor (GEF) that promotes dissociation of GDP from RAS protein in the brain in response to Ca^2+^ influx, muscarinic receptor signalling, or activation of the G protein beta-gamma subunit, and appears crucial for long-term memory formation in mouse model systems(Drake *et al.*, 2011, Barman *et al.*, 2014, Manyes *et al.*, 2018). Atxn10 (Ataxin 10) is a cytoplasmic protein required for neuron survival, differentiation and neuritogenesis via activation of mitogen-activated protein kinase cascade(Marz *et al.*, 2004, Waragai *et al.*, 2006). Lastly, Nedd4(Neuronal precursor cell-expressed developmentally downregulated 4) is an E3 ubiquitin ligase enzyme that promotes endocytosis and proteasomal degradation of various ion channels and membrane transporters, thereby contributing to the formation of neuronal dendrites, neuromuscular junctions, cranial neural crest cells, motor neurons, and axons(reviewed(Donovan and Poronnik, 2013)). Importantly, the critical roles played by Nedd4 is not limited to neurological functions alone, as it is also been implicated in tumorigenesis(Bergeron *et al.*, 2010, Zou *et al.*, 2015, Yang *et al.*, 2018). The potential applications of pSIVOM therefore extend beyond the study of neurodegenerative pathologies and could also be used to shed light on the pathogenesis of various cancers.

Our study presents an optimised protocol for studying protein dynamics *in vivo* that allows the molecular mechanisms underlying various pathologies to be investigated in a cost-effective and time-efficient manner. In the current report, we applied this approach to determine relative rates of protein turnover in different regions of the mouse brain, thus paving the way for future studies of how these dynamics are influenced by site-specific stimuli. It will be possible to use pSIVOM by altering feeding time points and sampling specific tissue regions to generate a highly detailed picture of proteomic regulation in the brain. Improving our understanding of protein physiology in the healthy brain will subsequently lead to advances in our knowledge of the mechanisms underpin dementia disorders. For example, pSIVOM could potentially be used to perform direct comparisons of different age groups of male and female mice to help clarify why the clinicopathologic features of dementia vary between genders in human patients, eventually leading to tailored therapies for different cohorts.

## Materials and Methods

### Animal housing and *in vivo* protein labelling

C57BL6/J mice aged 7-8 weeks were obtained from *InVivos* and housed in groups of three animals per cage. After 2-3 week adaptation to their new environment with provision of normal mouse chow (Altromin), the mice were fasted for 16h and transferred onto scaled SILAC diet (^13^C L-Lysine, SILANTET) with sterilized drinking water provided *ad libitum* for 2 days. Food intake, drinking water consumption, and body weight were monitored daily. All the experimental protocols were performed in accordance with the guidelines established by NTU Institutional Animal Care and Use Committee (NTU-IACUC), and all the methods were approved by the committee (IACUC protocol # ARF-SBS/NIE-A18018)

### Brain tissue processing

Mice were sacrificed with CO_2_ and immediately follow by cardiac puncture blood collection before brain tissues were excised, dissected, and immediately snap-frozen in liquid nitrogen or on dry ice. For total protein extraction, brain tissues were disaggregated using a liquid nitrogen cooled pulverizer (BioSpec) and 100mg of the resultant sample was re-suspended in 100mM ammonium bicarbonate (ABB, Sigma) lysing buffer containing 2% SDS together with protease inhibitor cocktail (Merck). The tissue suspension was then further homogenized using 1mm magnetic beads (Next Advance) in a bullet blender homogenizer (BioFrontier Technology) under high intensity at 4°C for 5min. The tissue homogenates were subsequently centrifuged at 10,000 x g, 4°C for 10min and the supernatants collected. Further rounds of homogenization were performed as required until no visible pellet remained. The collected supernatants were then combined, quantified and processed for in-gel digestion or precipitated with acetone (Fisher Chemical) prior to in-solution digestion. Acetone precipitation was perform for 4h at −20°C, follow by centrifugation at 5000xg, 4°C for 5min. The pellets were then air-dried and resolved in 100mM ABB buffer containing 8M urea and protease inhibitor cocktail.

### In-gel tryptic digestion

A total of 200μg brain protein per mouse was separated on 10% SDS-PAGE gels and the lanes were then cut into 8 separate bands and subjected to in-gel digestion. Each gel bands were further cut into approximately 1-2mm^2^ pieces and washed several times with 25mM ABB follow by 25mM ABB containing 50% acetonitrile (ACN, Fisher Chemical) until gel pieces are completely destain. The destained gel pieces were dehydrate with ACN and speedVac for 5-10mins. These gels pieces were reduced with freshly prepare 10mM Dithiothreitol(DTT, Sigma), 25mM ABB for 1hr at 60°C and then alkylated with 55mM Iodocetamide (IAA, Sigma), 25mM ABB in the dark at room temperature for 1hr. Gel pieces were dehydrated using ACN and subjected to overnight digestion with sequencing grade modified trypsin(Promega) at 37°C. Peptides were extracted with 50% ACN, 5% acetic acid (Merck), and dried using SpeedVac (Eppendorf), and then stored at −20°C until use.

### In-solution tryptic digestion and HPLC fractionation

A total mass of 600μg brain protein was subjected to reduction with 10mM Tris(2-carboxyethyl)phosphine(TCEP, Sigma) at 30°C for 2hrs and then alkylation with 20mM IAA containing 100mM ABB in the dark at room temperature for 30mins. Sequencing grade modified trypsin was added immediately and incubate at 37°C overnight. The tryptic peptides were then desalted with Sep-Pak C18 cartridge (Waters) and dried in a SpeedVac. Peptides were then dissolved with Buffer A (0.02% NH4OH) and subjected to high pH reversed-phase liquid chromatography fractionation with Buffer B (0.02% NH4OH, 80% ACN) on a C18 column (4.6×200mm, 5um, 300 Å, Waters, USA) at a flow rate of 1.0ml/min using a HPLC. The established 60-min gradient is set as 97% buffer A for 3min, 3-10% buffer B for 2min, 10-5% buffer B for 40min, 35-70% buffer B for 5min and 100% buffer B for 10min at 1ml/min flow rate. A total of 60 individual fractions were collected and then combined into 15 separate pools according to concatenation order. All fractions were subsequently SpeedVac dried and stored at −20°C until use.

### LC-MS/MS analysis

Four independent biological replicates were performed; brain samples 1-3 were prepared by in-solution digestion and HPLC separation into 15 fractions, while brain sample 4 was instead separated into 8 individual fractions by SDS-PAGE prior to in-gel digestion (Fig. 1). Tryptic peptides were re-suspended in 0.1% formic acid (FA, Fisher Chemical) as reserved volume, and each fraction was injected 3 times (TR1, TR2, and TR3 for in-solution digested samples) or twice (in-gel digestion sample) as technical replicates in LC-MS/MS. The peptides were separated and analyzed on a Dionex Ultimate 3000 RSLCnano system coupled to a Q-Exactive (Thermo Fisher). Approximately 2μg peptide from each fraction was injected into an Acclaim peptide trap column (Thermo Fisher) via the Dionex RSLCnano autosampler. Peptides were separated in a Dionex EASY-Spray 75μm × 10cm column packed with PepMap C18 3μm, 100 Å (PepMap® C18) at 35°C. Flow rate was maintained at 300nL/min. Mobile phase A (0.1% FA in 5% acetonitrile) and mobile phase B (0.1% FA in 90% acetonitrile) were used to establish a 60min gradient. Peptides were then analyzed on a Q-Exactive with EASY nanospray source (Thermo Fisher) at an electrospray potential of 1.5kV. A full MS scan (350–1600 m/z range) was acquired at a resolution of 70,000 at m/z 200 and a maximum ion accumulation time of 100ms. Dynamic exclusion was set as 15s. The resolution of the HCD spectra was set to 35,000 at m/z 200. The AGC settings of the full MS scan and the MS2 scan were 3E6 and 2E5, respectively. The 10 most intense ions above the 2000 count threshold were selected for fragmentation in HCD with a maximum ion accumulation time of 120ms. An isolation width of 2 was used for MS2. Single and unassigned charged ions were excluded from MS/MS. For HCD, the normalized collision energy was set to 28%. Underfill ratio was defined as 0.2%.

**Fig. 1.**
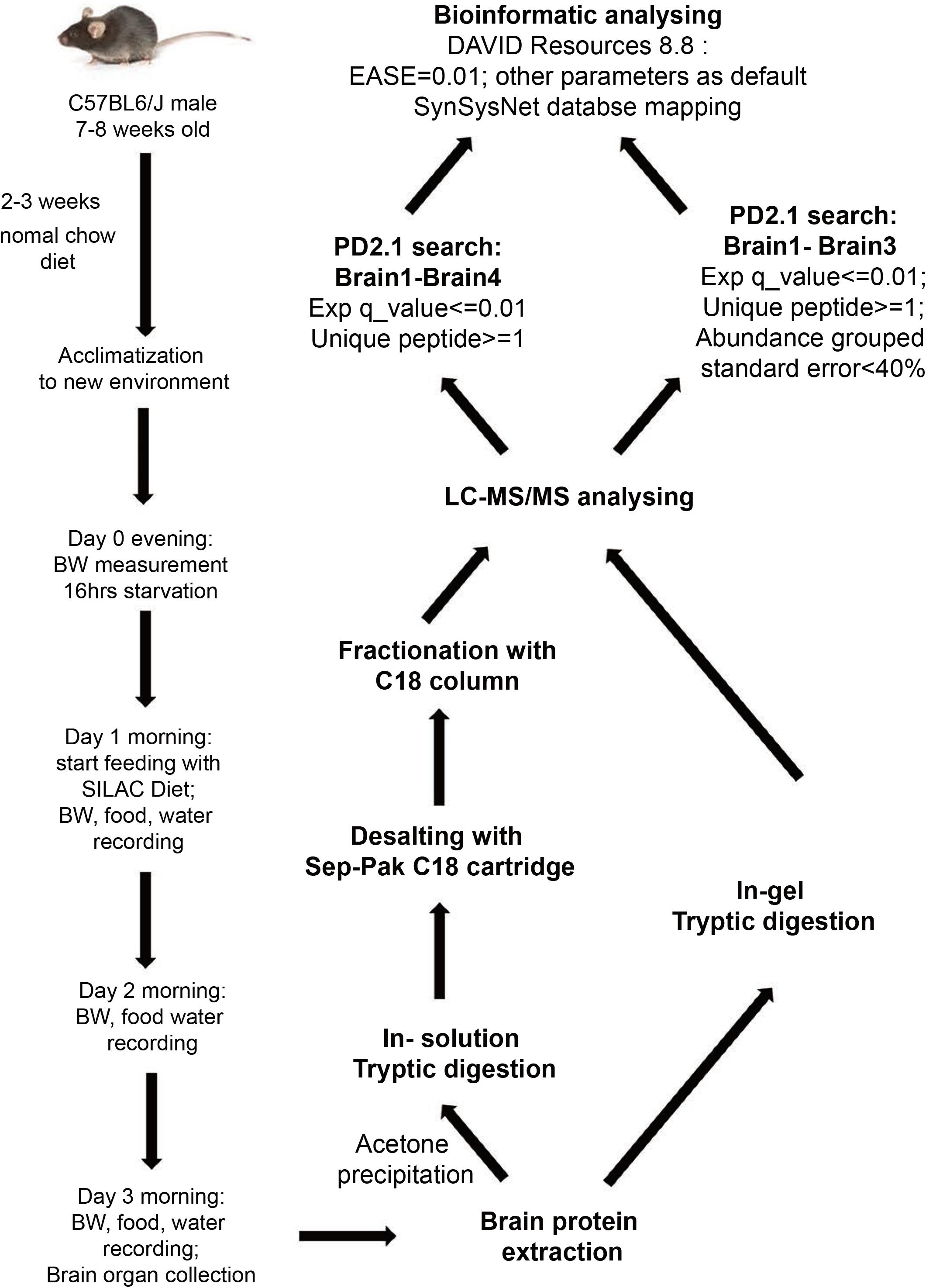
Workflow for *in-vivo* pulsed-SILAC labelling of mouse brain and subsequent proteomic analysis. Male C57BL6/J mice aged 7-8 weeks were acclimatised to their environment for 2-3 weeks on normal mouse chow diet containing ^12^C L-Lysine. The animals were then fasted for 16h overnight before switching to a ^13^C-L-Lysine-labelled SILAC diet for 2 days duration. Mice were then euthanized and the brain tissues were excised for protein extraction and mass spectrometry analysis. Pulsed-SILAC proteome analysis then allowed identification of newly synthesized proteins (^13^C L-Lysine-labelled) within specific regions of the mouse brain.

**Fig. 2.**
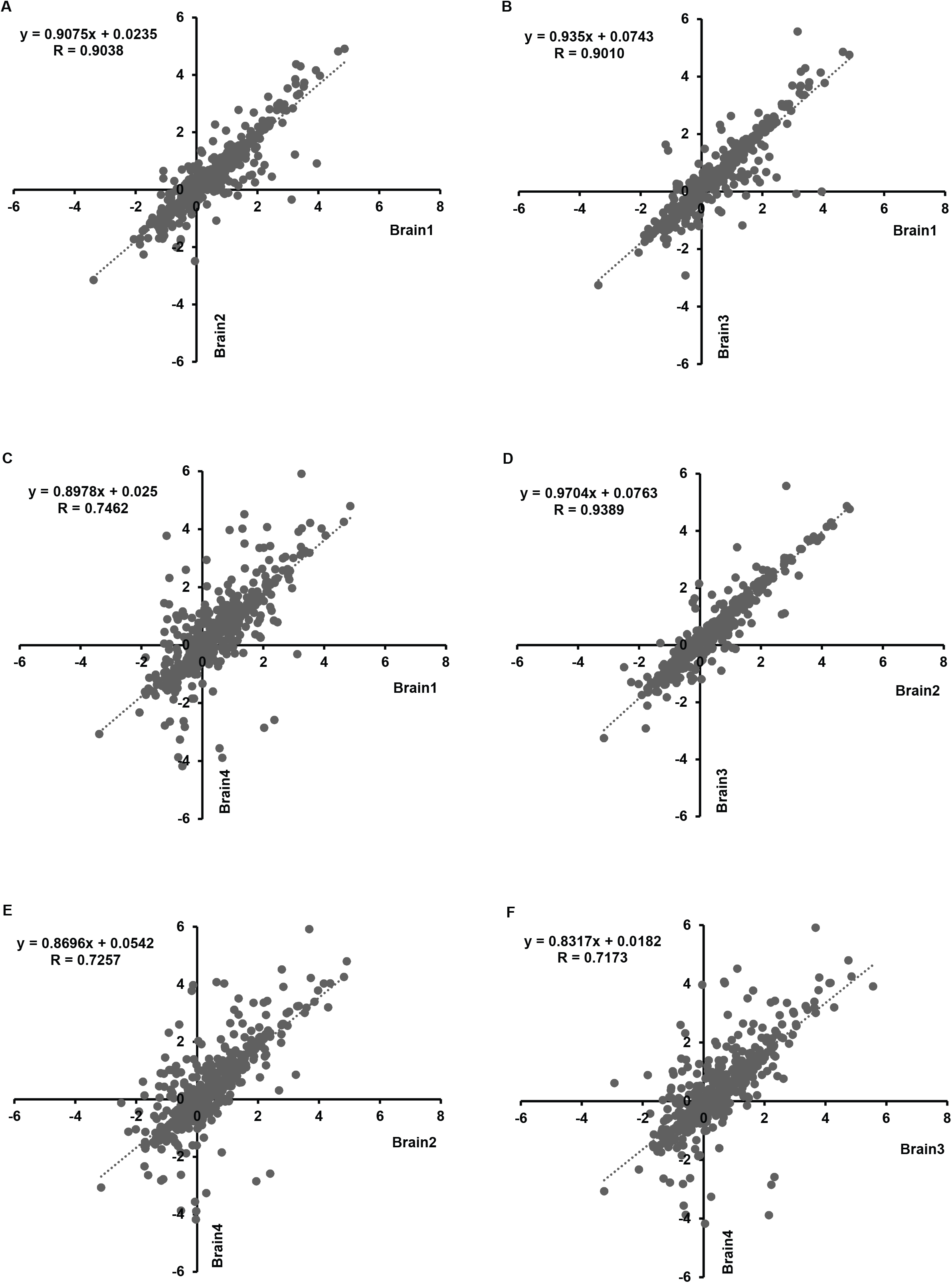
Correlation of 432 ^13^C-L-Lysine-labelled brain protein groups detected between biological samples indicating the high reproducibility of the proteomic data. X-axis v.s. Y-axis : (A) Brain1 v.s. Brain2; (B) Brain1 v.s. Brain3 ; (C) Brain1 v.s. Brain4; (D) Brain2 v.s. Brain3; (E) Brain2 v.s. Brain4; (F) Brain3 v.s. Brain4. Average heavy/light abundance ratio [Log2] values of each biological samples were used for correlation calculation and scatter plots plotting.

**Fig. 3.**
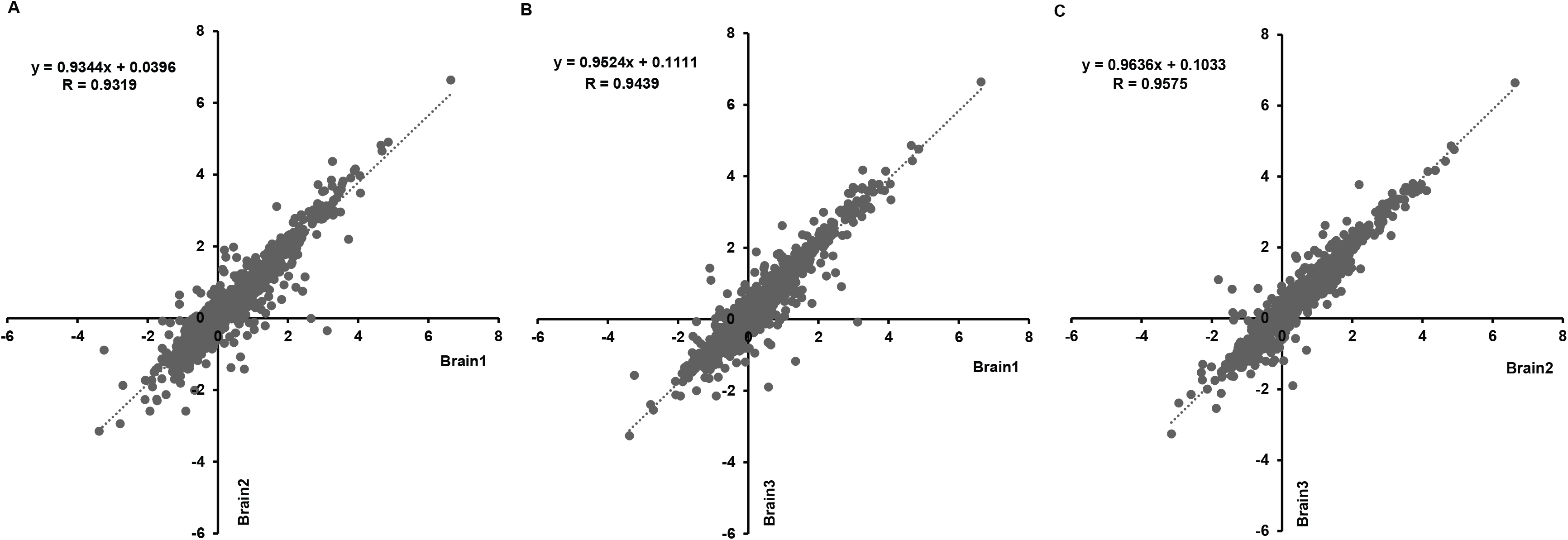
Brain protein groups identified with high confidence displayed strong correlation in the proteomic datasets generated by biological samples 1, 2 and 3. X-axis v.s. Y-axis: Brain1 v.s. Brain2; (B) Brain1 v.s. Brain3; (C) Brain2 v.s. Brain3. n=823, average heavy/light abundance ratio [Log2] value was used for correlation calculation and scatter plots plotting.

**Fig. 4.**
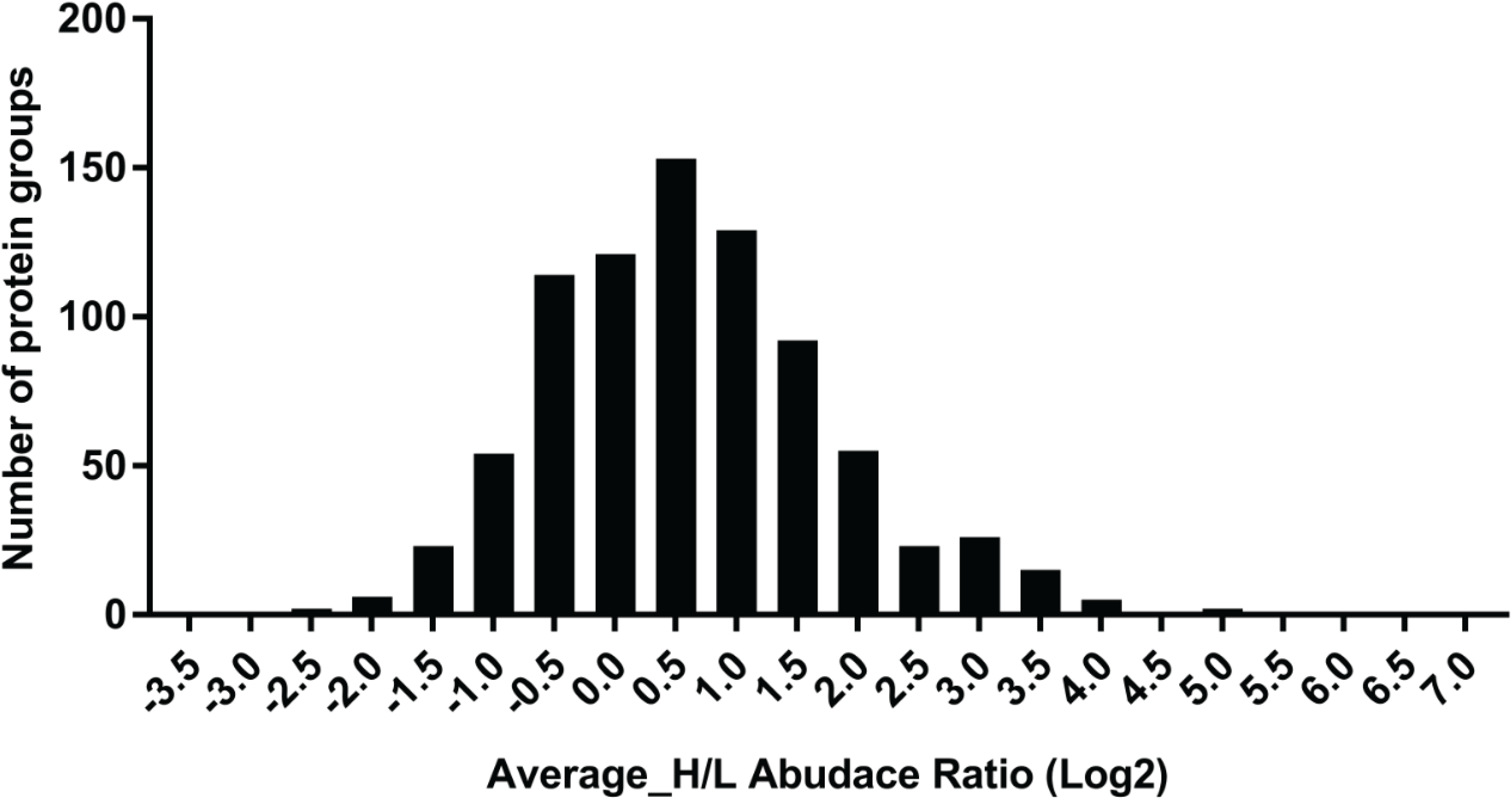
Distribution histogram on selected 823 protein groups. Protein average abundance (H/L ratio [Log2]) in selected 823 protein groups, represented as a distribution histogram for highly confident active translating proteomic datasets in Brain1, Brain2 and Brain3.

**Fig. 5.** Corpora quadrigemina and visual cortex are the most active translation region in brain. Different brain function related regions display variable newly synthesized protein abundance (average H/L ratio [Log2]). Based on the mean values, Corpora quadrigemina and visual cortex are most active regions in brain.

### Mass spectrometric Data analyzing

Raw data files from 11 replicates (three injections per biological replicates 1-3, plus two injections for biological sample 4) were analyzed as four independent experiments using Proteome Discoverer (PD) v2.1 (ThermoScientific) with the Uniprot mouse protein database (downloaded on 16 March 2017, 91089 sequences, 38788886 residues) using designed workflow. Briefly, this workflow includes six processing nodes numbered from 0 to 6. Node 0 named “Spectrum Files” allows selecting raw files, Node 1 labeled as “Spectrum Selector” extracts, deisotopes and deconvolutes the spectra within a retention time window and precursor ion mass window. Node 2 selected search engine SequestHT and Node 3 used Mascot with database search parameters. The parameters set were enzyme: trypsin, maximum miss cleavage:2, minimum peptide length:6, maximum peptide length: 6, maximum peptide length: 144, precursor mass tolerance: 10 ppm, fragment mass tolerance: 0.02 Da, modification groups (from quan method): SILAC K=6[MD], dynamic modification: deamidation of N and Q, Oxidation(M), Static modification: Carbamidomethyl(C), Node 4 called as “Percolator” where target FDR(strict) was set as 0.01, target FDR(relaxed) was set as 0.05. Node 5 labeled as “Event Detector”, Mass precursor set as 4ppm, S/N Threshold set as 1, Node 6 labeled as “Precursor ions Quantifier”, where RT Tolerance of Isotope pattern multiplets(min):0.2 and Single peak\Missing channels allowed:1. The obtained peptide/protein list was further analyzed with the designed consensus workflow with PD2.1. In short, total nine nodes, Node 0 named as MSF Files which the “spectra to store” set as both identified or quantified, “merging of identified” is set as gloabally by search engine type, and the “reported FASTA Title lines: Best match, Node 1 is “PSM Grouper” which “peptide group modification the site probability Threshold set as 75 and ‘modification sites” show only best position, Node 2 is “Peptide and Protein filter”, “Validation mode set as “automatic and the target FDR for both PSMs and Peptides were set as “strict:0.01 and “Relaxed”: 0.05, Node 3 is “Peptide and protein filter” peptide confidence: high, minimum peptide length: 6, minimum peptide sequence: 1, Node 4 was “Protein scorer” which including Node 5 that named as “Protein grouping” : the strict parsimorny principle was applied, Node 6 labelled as “ Protein FDR Validation” : 0.01 as strick and 0.05 as relexed, Node 7 was named as “Peptide in protein annotation” and Node 8 was “Peptide and Protein quantifier”: use Unique+Razor peptide, consider protein groups for peptides unique, Normalized mode: total peptides, Scaling mode: On channels average(per file), Report Quantification: Reproter Abundance Based on: intensitys, Co-isolation Threshold: 50, Average reporter S/N Threshold: 100

### Bioinformatic analysis

Bioinformatics analysis was performed using DAVID Resources 6.8 (EASE=0.01, others as default setting) (Huang da *et al.*, 2009). The synaptic protein/gene database was downloaded from the SynSysNet website (von Eichborn *et al.*, 2013).

### Statistical analysis

Each set of PD2.1 search peptide/protein list was exported to Microsoft Excel and then subjected to cut-off filtering according to the following parameters; Exp q_value≤ 0.01, Unique peptide≥1, Abundances grouped standard error in percent Light and Heavy <40% (except for brain sample 4). The statistical analysis was performed by PD2.1 default setting. Pearson correlation coefficients (default Excel Correl function) were used to measure the strength of the relationship between biological and technical replicates; correlation coefficient values >+0.8 is consider as significant positive correlated. Standard Deviation (SD) was used to measures the variance between biological triplicates of interested proteins. The distribution Figures were generate and analysed with GraphPad Prism 7.

## Data availability

LC-MS/MS raw data from the 11 replicates and results for protein and peptide identification and quantification from PD2.1 have been deposited to the ProteomeXchange Consortium via the PRIDE(Perez-Riverol *et al.*, 2019) partner repository with the dataset identifier PXD013502. The raw data and search results can be accessed using the following login to the PRIDE data depository.

Username

Password: hdkeFqIl

## Supplementary

### Supplementary Figure Legends

**Supplementary Fig. S1. Record of body weight for individual mice (A) and average daily SILAC food intake per animal (B).** 10 weeks old C57BL6 mice were caged as three mice per cage. Their body weight, food and water intake were measured and recorded as indicated.

**Supplementary Fig. S2. Selected newly synthesized protein groups identified with high confidence (displayed together with correlation between technical triplicates)**; (A) Brain1_TR1 correlation with Brain1_TR2. (B) Brain1_TR1 correlation with Brain1_TR3. (C) Brain1_TR2 correlation with Brain1_TR2. (D) Brain2_TR1 correlation with Brain2_TR2. (E) Brain2_TR1 correlation with Brain2_TR3. (F) Brain2_TR2 correlation with Brain2_TR3. (G) Brain3_TR1 correlation with Brain3_TR2. (H) Brain3_TR1 correlation with Brain3_TR3. (I) Brain3_TR2 correlation with Brain3_TR3. n=823; heavy/light abundance ratio (Log2) values were used for correlation calculation and scatter plots plotting.

**Supplementary Table S1. Record of mouse body weight and average SILAC diet consumption per animal.**

**Supplementary Table S2. Brain protein groups identified with high confidence within the ^13^C-L-Lysine-labelled fraction from any 3 biological sample**s.

**Supplementary Table S3. Brain protein groups identified with high confidence within the ^13^C-L-Lysine-labelled fraction from all samples analysed.**

**Supplementary Table S4. Selected newly synthesized protein groups identified with high confidence (displayed together with correlation between technical triplicates).**

