## Supplementary figures and images for "Dynamics of murine brain protein synthesis *in vivo* identify the hippocampus, cortex and cerebellum as highly active metabolic sites"

### Supplementary Figure S1

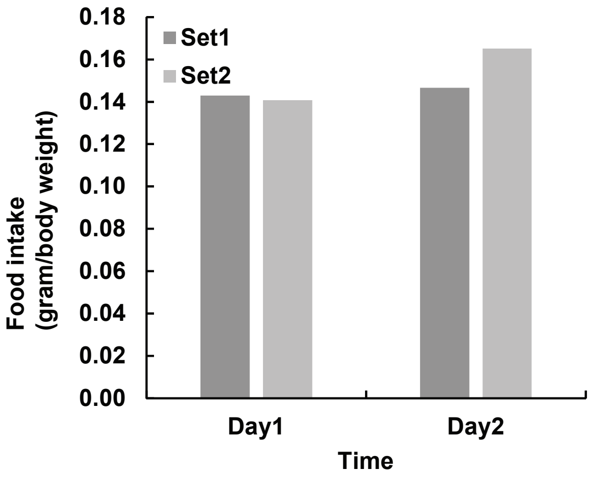

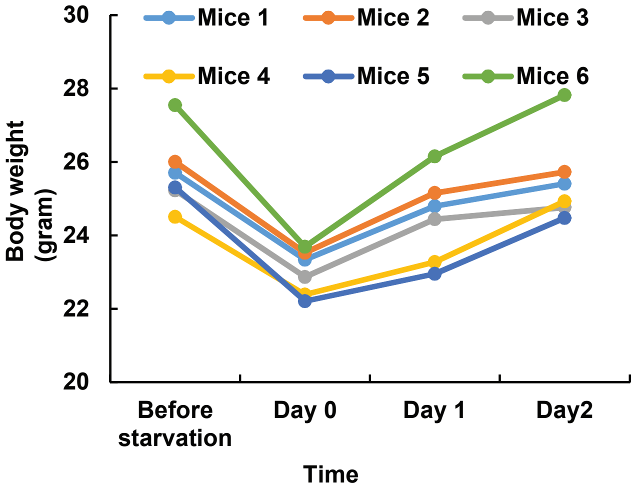

### Supplementary Figure S2

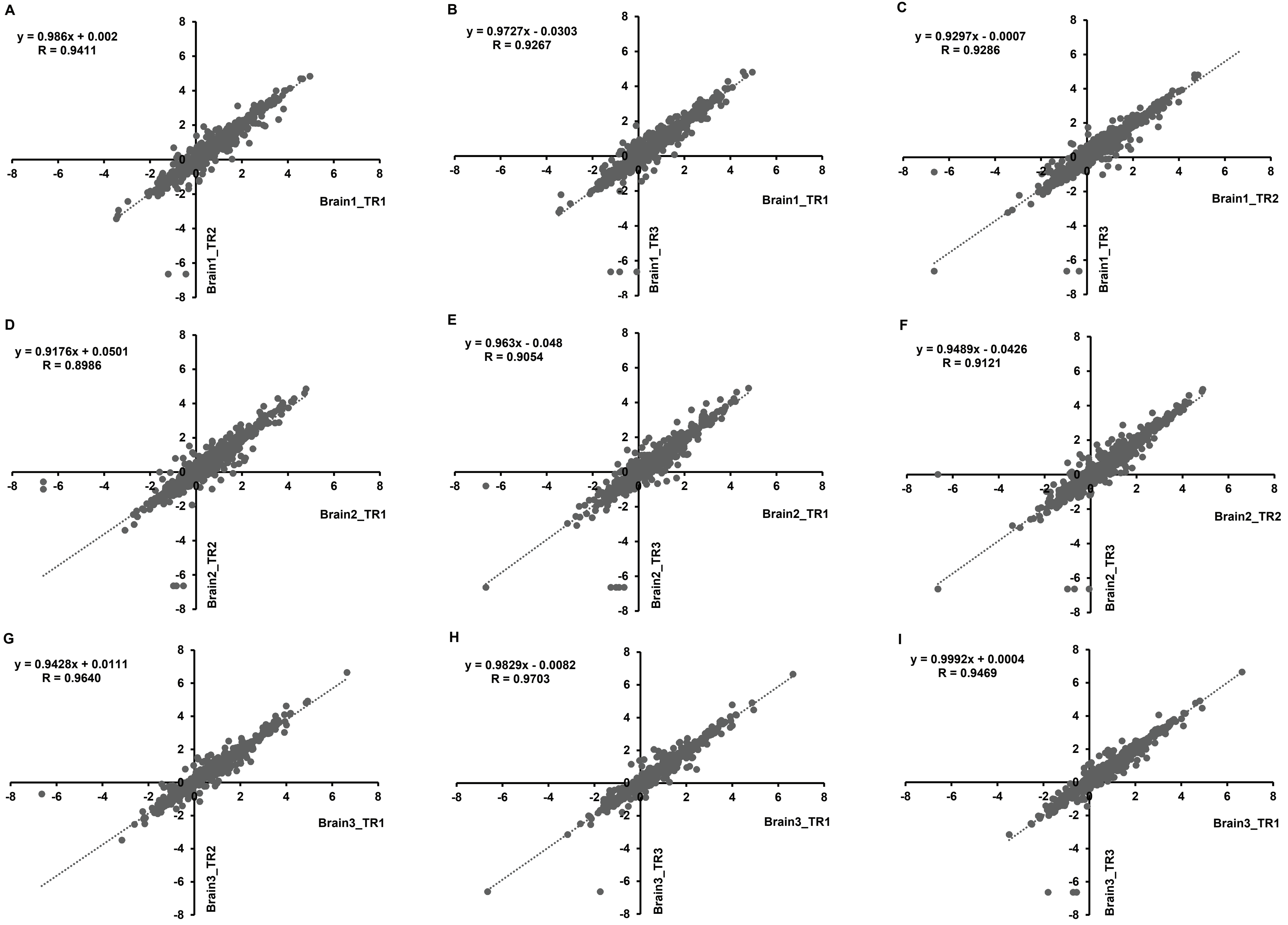
